# Feature-based Quality Assessment of Middle Cerebral Artery Occlusion Using 18F-fluorodeoxyglucose Positron Emission Tomography

**DOI:** 10.1101/2020.12.28.424532

**Authors:** Wuxian He, Hongtu Tang, Jia Li, Chenze Hou, Xiaoyan Shen, Chenrui Li, Huafeng Liu, Weichuan Yu

**Author notes:** Contributed equally (Wuxian He, Hongtu Tang, Jia Li). Corresponding author: Email addresses (Huafeng Liu), (Weichuan Yu).

## Abstract

**Background:** The association between brain metabolic change and ischemic stroke has attracted a lot of attention in the research community. 18F-fluorodeoxyglucose (FDG) positron emission tomography (PET) imaging is widely used to measure the metabolism. In experiments, ischemic stroke is usually induced through middle cerebral artery occlusion (MCAO), and quality assessment of this procedure is of vital importance. However, an assessment method based on FDG PET images is still lacking. Herein, we propose an image feature-based protocol to assess the quality of the procedure.

**Methods:** We performed permanent MCAO to a total of 161 Sprague-Dawley rats. FDG micro-PET images were acquired both before and after the MCAO procedure. Triphenyl tetrazolium chloride (TTC) staining was also conducted to obtain ground truth of the infarct volume. After preprocessing of the PET images, a combination of 3D scale invariant feature transform (SIFT) and support vector machine (SVM) was applied to extract features and train a classifier that can assess the quality of the MCAO procedure.

**Results:** 106 rats and 212 images were used as training data to construct the classification model. The SVM classifier achieved over 98% accuracy in cross validation. 10 rats with TTC results showing infarction in the ipsilateral brain region served as validation data. Their images were tested by the classifier and all of them were categorized into the correct group. Finally, the remaining 45 rats from a separate experiment were treated as independent test data. The prediction accuracy for these 90 images reached the level of 91%. An online interface was constructed for users to upload their images and obtain the assessment results.

**Conclusion:** This feature-based protocol provides a convenient, accurate and reliable tool to assess the quality of the MCAO procedure in FDG PET study.

## 1. Introduction

Stroke is one of the most serious brain diseases and the second leading cause of death around the world [1]. During an acute stroke, the brain will undergo dysfunction and certain areas will be damaged. To study the stroke mechanism and treatment effects, animal subjects have been extensively used. Among animal subjects, rats are very popular because of their low cost and similar physiology to humans. To induce focal cerebral ischemia in rat subjects, middle cerebral artery occlusion (MCAO) is the most commonly used procedure [2]. After the occlusion, assessments need to be performed to ensure the success of the MCAO before follow-up experiments can be carried out. Traditionally, behavior tests [3], and triphenyl tetrazolium chloride (TTC) staining [4], were used to assess the MCAO quality. However, the behavior test is subjective, and the judgments vary among different examiners [5]. Although TTC staining can show the infarct volume and provide the ground truth of the MCAO quality, it needs to sacrifice rats. Consequently, follow-up experiments become infeasible. In recent years, more assessment methods based on imaging modalities have gained popularity due to their objective and non-invasive nature. Laser Doppler flowmetry (LDF) [6, 7] can monitor the blood flow in brain areas connected to the occluded vessels in real time. Studies have shown significant correlation between the LDF change and behavior scores or infarct volume [8]. However, limitations exist for this technique such as spatial heterogeneity due to tissue perfusion and motion [9]. Functional magnetic resonance imaging (fMRI) measures the blood-oxygen-level dependent (BOLD) contrast signal, which is associated with the brain oxygen metabolism [10]. However, it is quite sensitive to movements. Consequently, the head motion of rats during the scan session may generate undesired blurring effect [11, 12]. Positron emission tomography (PET) is another popular functional imaging technique that uses radioactive substances to measure the *in vivo* information such as physiological activities and metabolism [13, 14].

There are two common types of radioactive tracers used in PET study of stroke: 15O-H_2_O and 18F-fluorodeoxyglucose (FDG). PET imaging with 15O-H_2_O provides an indirect measure of cerebral blood flow (CBF) [15]. Decreased 15O-H_2_O concentration can be recorded in rats after MCAO, which associates with reduced rate of perfusion [16]. However, the half-life of the oxygen-15 isotope is only about 2 minutes, which requires onsite production using a cyclotron. Few laboratories can meet this requirement on apparatus and thus the FDG radiotracer becomes more popular for research experiments. PET imaging with 18F-FDG studies how brain metabolism varies with cerebral ischemia and infarction [17]. Because the regional glucose level is directly related to brain metabolism, the FDG concentration can accurately reveal the metabolic change. Reduced FDG concentration in the injured areas indicates diminished level of neuronal activity. Analysis approaches have been proposed to assess the MCAO quality based on PET images, among which the statistical parametric mapping (SPM) gains most credit [18]. The SPM uses general linear model (GLM) to represent data and conduct hypothesis testing (*t* tests or *F* tests) to identify voxels/regions that show significant change before and after stroke [19]. PET studies using 15O-H_2_O have shown consistent results where significant decrease of radiotracer concentration (in terms of image intensities) were observed in affected areas, demonstrating reduced CBF [20]. PET studies using FDG, however, have a large variability. Some works reported decreased FDG uptake in brain regions corresponding to the ischemic core from hours to days [21], which are reasonable because reduction of CBF directly suppresses the glucose supply. However, there are works that showed alteration in spatiotemporal uptake of FDG [17, 22]. Such inconsistent results exist in the peri-ischemic regions, where either insignificant change or hyper-uptake of FDG were reported. The underlying reasons for the increased FDG uptake in the ipsilateral hemisphere can be attributed to the activation of glucose transporter (GLUT), inflammation or neuronal regeneration [17]. Therefore, significant results in FDG PET image analysis may not be consistently accessible using voxel-wise activation methods.

Performing SPM for FDG PET images also suffers from the problem of intensity normalization [23]. Since the dosage of injected radiotracer cannot be controlled to be identical for each rat, concentration calibration must be carried out. This requires dividing each voxel by the global mean intensity over the intracranial volume. However, bias and errors could arise due to various factors such as variations of scanning time, plasma glucose level and basal metabolic rate [24]. Hence, a fast, accurate and reliable approach to assessing the MCAO quality based on FDG PET scans is desirable.

Here, we adopt a 3D scale invariant feature transform (SIFT) algorithm [25] to extract features from the FDG microPET images. The original 2D SIFT [26] is a well-known feature detection approach that can extract local scale- and rotation-invariant features from images. There are mainly four steps for the SIFT procedure and they are briefly described in the Online Data Supplement. Then, a support vector machine (SVM) classification model is built from the features of labeled images, which can classify an unknown new image into either the healthy or MCAO category with high accuracy. Existing approaches for FDG PET image analysis require two scans per subject: one baseline and one taken after MCAO [27]. In practice, preparing the radiotracer and taking PET images twice for each rat are expensive and time consuming, especially when a large number of rats are involved in the experiment. Our protocol overcomes such a defect by building a model that contains feature information from both baseline and MCAO images. Consequently, only one scan after the MCAO is needed for the assessment. Moreover, since the feature extraction does not require calibration, this analysis pipeline also avoids potential errors introduced during the intensity normalization process.

In the following sections, we shall describe how to perform the image feature-based assessment, from preprocessing, through feature extraction, to model construction, and demonstrate the significance of the protocol.

## 2. Methods

### 2.1. Animals

All the animal experiments in this study were approved by the Animal Research Committee of School of Medicine, Zhejiang University. Two independent MCAO surgeries were performed following the same procedure [2], one with 116 rats and the other with 45 rats. The subjects involved in this study were adult female Sprague-Dawley rats with a weight of 180-280 g. During the experiment, each rat had free access to food and water, and was anesthetized using 400 mg/kg chloral hydrate before the scanning. The entire experimental procedure followed the guidelines for the care and use of Laboratory Animals published by the National Institute of Health. All the data are available from the website: http://yulab.ust.hk/MCAO/

### 2.2. Image data acquisition

All 3D FDG microPET images were acquired with a microPET R4 system (CTI Concorde Microsystems, LLC.) located at the Medical PET Center of Zhejiang University. For each rat in this study, one scan was taken for the resting state (baseline) before the MCAO procedure and one was taken after the MCAO procedure. Both scans were acquired after 30 minutes of 18F-FDG injection and each lasted for 15 minutes. The spatial resolution of the scanner was 1.9 mm full width at half maximum (FWHM) in the transverse plane and 1.88 mm FWHM in the axial plane. Additionally, 10 rats from the first experimental group were chosen randomly to have TTC staining. The results provided clear visualization of the tissue damage in the right brain, and photos were taken as a record for validation.

### 2.3. FDG PET template

To transform PET scans of distinct rats into a standard space, an anatomical template was necessary. Since there is no widely accepted FDG PET template for Sprague-Dawley rats, we constructed our own template from an existing MRI template [28] using SPM12 (Wellcome Department of Cognitive Neurology, London, UK). The detailed steps are described in the Online Data Supplement. The MRI template we used and the PET template we constructed are shown in Figure 1.

**Figure 1:**
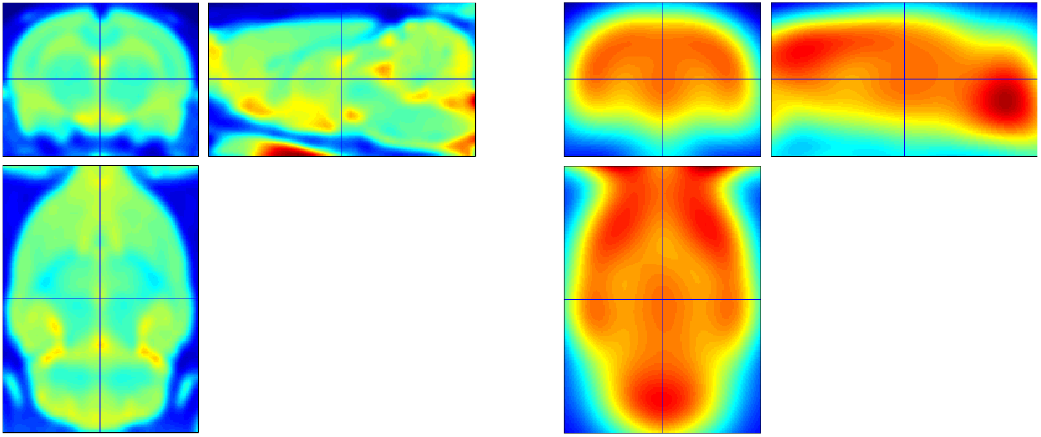
Templates used in image preprocessing. Left: an existing MRI template for Sprague-Dawley rats; Right: the FDG PET template for Sprague-Dawley rats that we constructed.

### 2.4. Preprocessing

The raw data were reconstructed using the maximum a posterior (MAP) algorithm [29]. Then, scans were preprocessed with the constructed template in SPM12. The pipeline of the preprocessing is shown in the Online Data Supplement. Figure 2 shows an example of a raw image and its preprocessed result.

**Figure 2:**
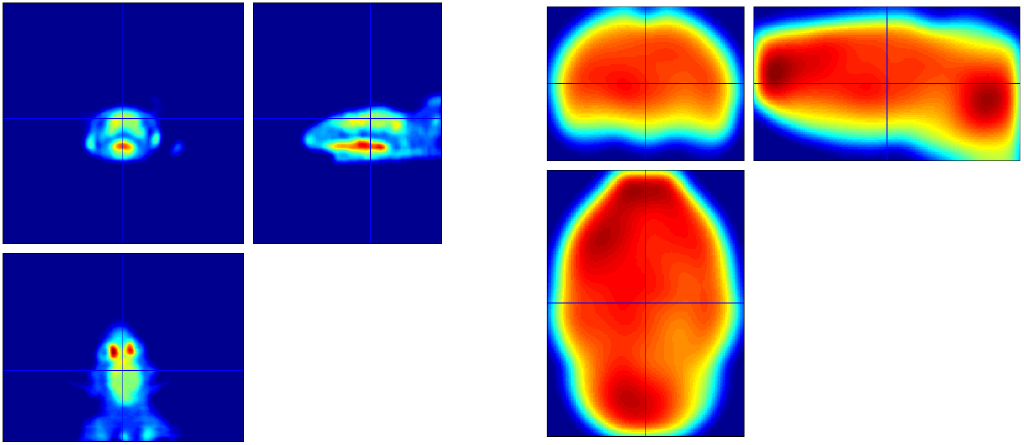
An example of a raw scan after reconstruction was shown on the left. Brain size were not unified, images were not aligned well, and unnecessary components such as eyes were superfluous. The final result after preprocessing on the right only contained intracranial volume in a standard template space.

### 2.5. Image features

After spatial preprocessing, image features were extracted using the 3D SIFT method, which returned a final descriptor as a vector in 768 length [25]. Since the images had already been registered, we used a set of fixed key points and omitted the key point localization step. Furthermore, because SIFT extracts local features from an image, we proposed to choose key points from brain areas that are possibly affected during a stroke, such as the neocortex or striatum [30]. The delineation of brain regions (one slice is shown in Figure 3) was based on the Waxholm atlas made by Papp et al [31]. Because the symptoms and injured areas resulting from a stroke are not totally understood, in order to be inclusive, after many trials we determined 25 key points from 13 brain regions, as shown in Table 1. Then the feature of each image was formed by concatenating the descriptors of all the 25 key points of it, which became a vector in 768 × 25 = 19200 dimensions. In Figure 3 and the Online Data Supplement, we provide results showing the 3D rendered atlas along with the location of the 25 key points. The 3D rendering was completed by using an open-source data visualization application called ParaView [32].

**Figure 3:**
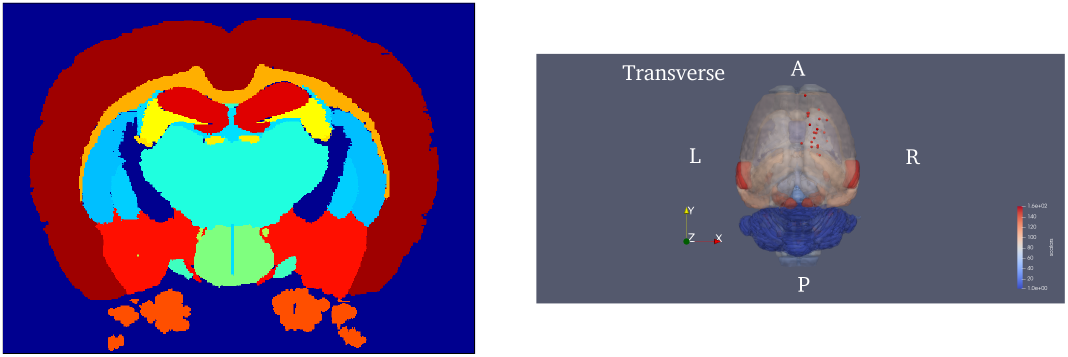
Left: a coronal slice of the Waxholm atlas showing brain region delineation; Right: a transverse view of the 3D rendered atlas, with the 25 key points (red dots) chosen from the ipsilateral hemisphere for feature extraction.

**Table 1:** The brain regions from which the 25 key points were chosen. Regions with more voxels and severe damage, such as neocortex and striatum, have multiple key points. The regions are listed in the same order according to the region index provided in the Waxholm Space atlas.

| Region name | Number of key points |
| --- | --- |
| Subthalamic nucleus | 1 |
| Alveus of the hippocampus | 1 |
| Striatum | 3 |
| Ventral hippocampal commissure | 1 |
| Thalamus | 2 |
| Fimbria of the hippocampus | 1 |
| Stria medullaris of the thalamus | 1 |
| Interpeduncular nucleus | 1 |
| Middle cerebellar peduncle | 1 |
| Neocortex | 10 |
| Postrhinal cortex | 1 |
| Perirhinal area 35 | 1 |
| Perirhinal area 36 | 1 |
| Total number: | 25 |

### 2.6. Classifier training

The calculations and statistical analysis were done in MATLAB R2018b (Natick, Massachusetts: The MathWorks Inc). The image features of all the training data were used to learn a linear SVM classification model. This model can be used to classify any new image into one of the two groups: normal (baseline) or MCAO (injured). The assessment of MCAO quality is based on the classification or prediction accuracy. Ten-fold cross validation was used to examine the performance of the model, and classification on the validation set and test set was also conducted to evaluate the performance.

### 2.7. Data availability statement

All the data and tools included in this paper are available for open access on the webpage: http://yulab.ust.hk/MCAO/. Furthermore, to facilitate the application of our method, we constructed an online interface that can automatically perform the image preprocessing, feature extraction and MCAO assessment. Users only need to do one manual step that roughly align their images to one example we provided before they upload. This is to ensure the accuracy of preprocessing results because we assume that different PET scans will have varying resolution and orientations.

## 3. Results

### 3.1. Key points and features

The brain regions from which we chose key points, along with the number of key points in each of them, are shown in Table 1. The neocortex was the largest region and contained voxels in a wide range. Thus, we chose the largest number of key points from it. Meanwhile, the striatum was the second largest region, and therefore we chose 3 key points from it. The decision to use these brain regions and key points was based on existing stroke studies [30, 33] and on our repeated experiments with different sets of key points.

The feature of each microPET image was obtained by concatenating the 25 SIFT descriptors into a single vector. As mentioned previously, we have 106 rats in the training set, 10 rats (with TTC ground truth) in the validation set and 45 rats (from an independent experiment) in the test set. Every rat was given two microPET scans, one before the MCAO surgery, which we call the normal image, and one after the surgery, which we call the MCAO image. Prior to training the SVM classification model, we performed principal component analysis (PCA) to the image features. As an unsupervised learning method, PCA can significantly reduce the feature dimensionality by projecting the data onto its principal vectors. This enables us to project the image features from a 19200-dimensional space to a 2-dimensional plane and visualize them. Figure 4 shows the PCA results of the two sets of image features, and both of them indicate a clear discrimination between normal and MCAO images. This label-free result suggests that the use of SIFT descriptors has promising accuracy to assess MCAO quality.

**Figure 4:**
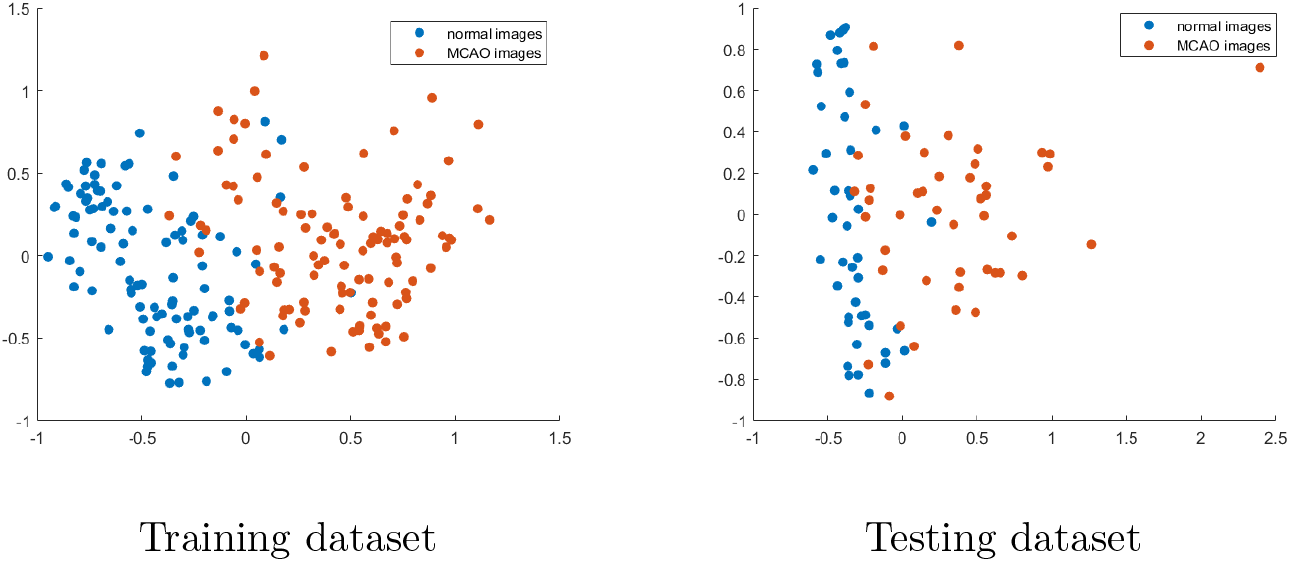
The PCA results of the two sets of SIFT features. Features in the original 19200dimensional space were projected to a 2-dimension plane. Left: images from the training set; Right: images from the test set.

### 3.2. Prediction accuracy

Although PCA returned significant results, its computation requires grouped data. Therefore, it cannot be applied to a single test image. For this reason, we turned to use supervised classification approaches. In order to make our protocol as simple as possible, we used basic and well-known classification methods. We chose the following traditional methods: K-nearest neighbor (KNN), decision tree (DT), linear discriminant analysis (LDA) and support vector machine (SVM) and tested their accuracy with cross validation [34]. The results shown in Figure 5 suggest that SVM is the optimal method with the highest accuracy. The classification accuracy with cross validation reached over 98% and the prediction accuracy for the test set was 91%. SVM was therefore adopted for the protocol.

**Figure 5:**
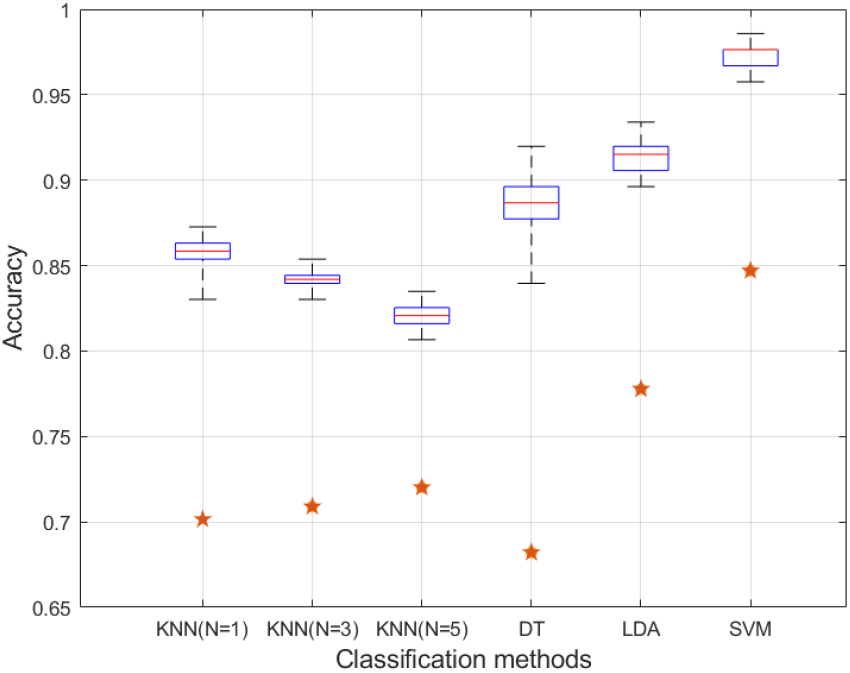
Box plot and scatter plot of accuracy for different classification methods. The box plot represents the 10-fold cross validation accuracy with different classification methods using the 106 rats from the training set. The orange scatter points were obtained from predicting the 45 rats in the independent test set using each trained model.

### 3.3. Validation with biomedical indicator

All the analyses so far treated the rats after MCAO as if their surgery were completed successfully. Although the MCAO procedures were carried out carefully, the results became less convincing if the ground truth of occlusion could not be validated through a direct measure. Therefore, we performed TTC staining on 10 randomly selected rats and the results all showed obvious brain infarction in the right half of the brain (one example is shown in Figure 6 and all staining photos are included in the Online Data Supplement). We then tested the 20 images (10 normal + 10 MCAO) with the SVM model trained from the 106 rats, and obtained 100% accuracy. All the TTC staining photos of the 10 rats are shown on our webpage.

**Figure 6:**
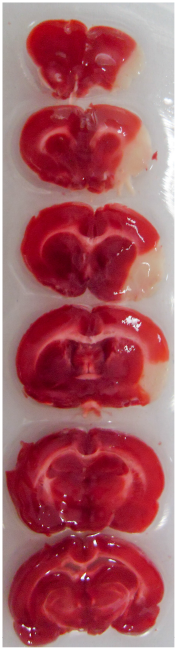
An example of the TTC staining result after 15 hours of occlusion.

## 4. Discussion

### 4.1. Comparison with intensity-based analysis

Our study was conducted using a large number of rats with microPET images acquired before and after MCAO. The size of our dataset was larger than that in most of the existing MCAO studies, and therefore we could investigate various methods and obtain reliable results for the assessment. Our protocol can also be applied to assess the MCAO quality using only a single image, and does not require the scan of the normal state before occlusion. This is a significant advantage over the current intensity-based assessment approaches such as SPM. The existing intensity-based methods all require two scans of the subject: one baseline image and one after MCAO, and significant findings can be made only when a clear intensity change between the two images is observed in an injured area [15, 35, 36].

Moreover, we found that signal changes among different rats were not consistent after we investigated the intensity histograms of subtracted scans. In the FDG PET study of the MCAO procedure, the intensities will directly indicate the uptake of glucose, and assessment of the MCAO is feasible if a significant intensity decrease is observed. However, the intensity value of a PET scan is also related to the radiotracer concentration. Because the concentration cannot be maintained at the same level for each injection, the intensity needs to be normalized by the average intensity value among all the intracerebral voxels for each rat [23]. Such intensity normalization steps produce the standardized uptake value (SUV), and the SUV change in two significant voxels and regions before and after MCAO from our dataset is shown in Figure 7.

**Figure 7:**
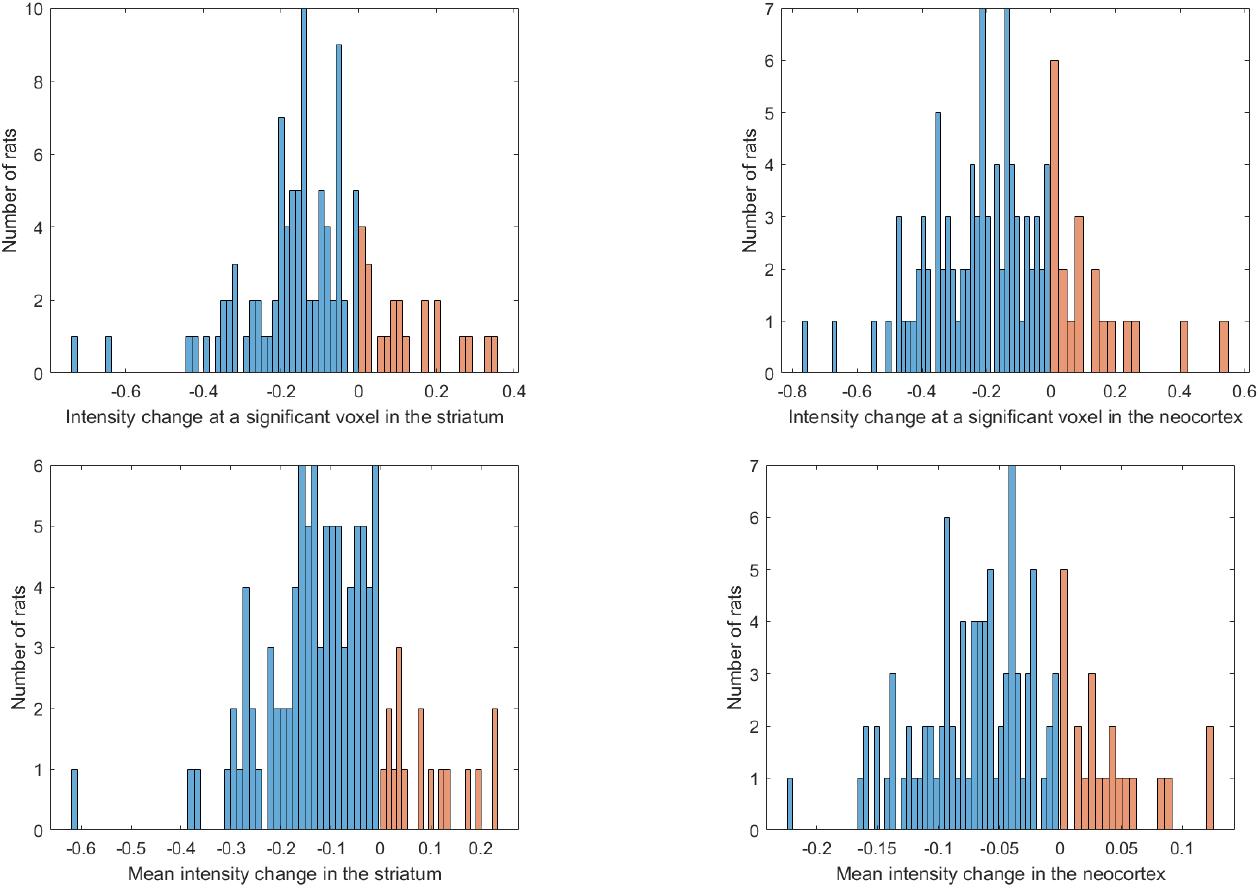
Histograms of the FDG uptake (normalized intensity) change between the normal and MCAO images (MCAO - normal) for all rats in the training set from two regions: the striatum (left) and the neocortex (right). The top two histograms belong to a single significant voxel from 2-sample *t*-test. The bottom two histograms belong to the mean intensities in the two regions. Orange bins represent the subjects with increased FDG uptake in the ipsilateral part after MCAO.

Figure 7 shows that the SUV change on two significant regions has nonuniform results. About one quarter of the total rats had increased FDG intake after MCAO even though we performed intensity normalization. As discussed previously, this phenomenon could have two explanations. One is the heterogeneous distribution of glucose in the peri-ischemic areas [22] and the other is the error from intensity normalization [24]. For normalization, the mean intensity in the intracranial part is largely determined by the healthy regions since the injured area will have diminished FDG uptake. If two rats have significantly different amounts of FDG injected, the overall intensity in the whole brain will vary significantly, and thus the denominator becomes a dominating factor [37]. In this way, the normalized intensity in the injured brain may even increase after MCAO if less FDG is injected before the scanning session. Using our feature-based protocol, we could overcome this problem and bring higher accuracy (Table 2). The heterogeneous uptake of FDG in the peri-ischemic regions can also be well included when we have more subjects in the training dataset. At the same time, taking only one scan per rat could save time and cost for researchers.

**Table 2:**
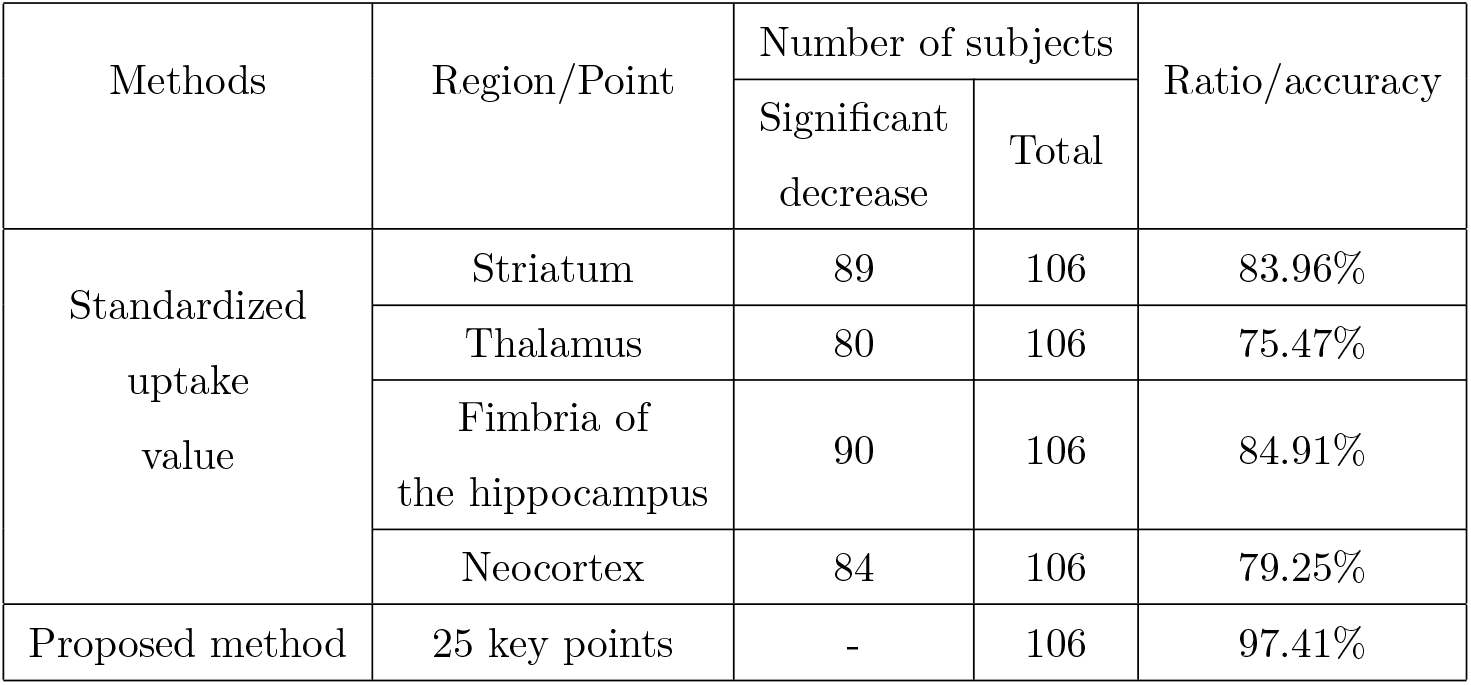
Comparison of results between using SUV and our proposed classification method. SUV change was calculated in 4 regions of interest: striatum, thalamus, fimbria of the hippocampus and neocortex from the training dataset. The number of rats that showed significant intensity decrease (p< 0.05, one-sided t-test) were counted and divided by the total number. The ratio was compared with the classification accuracy using our feature-based method.

### 4.2. Robustness of the FDG PET template

We provided a ready-to-use FDG PET template of Sprague-Dawley rats for raw image preprocessing. This template could be useful to researchers who desire a FDG PET template but lack data to construct one themselves. To ensure the 20 baseline images for the construction were able to build a representative and universal template, we studied multiple choices of the baseline images and verified the template quality. Since we had two independent datasets, we tested 7 different combinations of images:

1. 20 baseline images with good initial alignments carefully chosen from the training set.
2. 20 baseline images randomly chosen from the training set.
3. 10 baseline images randomly chosen from the training set and 10 baseline images from the test set.
4. 20 baseline images randomly chosen from the test set.
5. 20 baseline images with a high intensity level (12.92 17.44) from the training set.
6. 20 baseline images with a low intensity level (1.848 2.786) from the training set.
7. 10 baseline images with a high intensity level and 10 with a low intensity level from the training set.

We examined the quality of the constructed templates based on SVM accuracy. The results shown in Table 3 indicate that the choice of baseline images had a minor influence on the template quality and preprocessing steps. Even when we selected the baseline images only from one dataset and performed preprocessing and analysis on the other dataset, the result showed very good consistency. Therefore, the final template we fixed was the one from (1) above using 20 optimal baseline images from the training set.

**Table 3:**
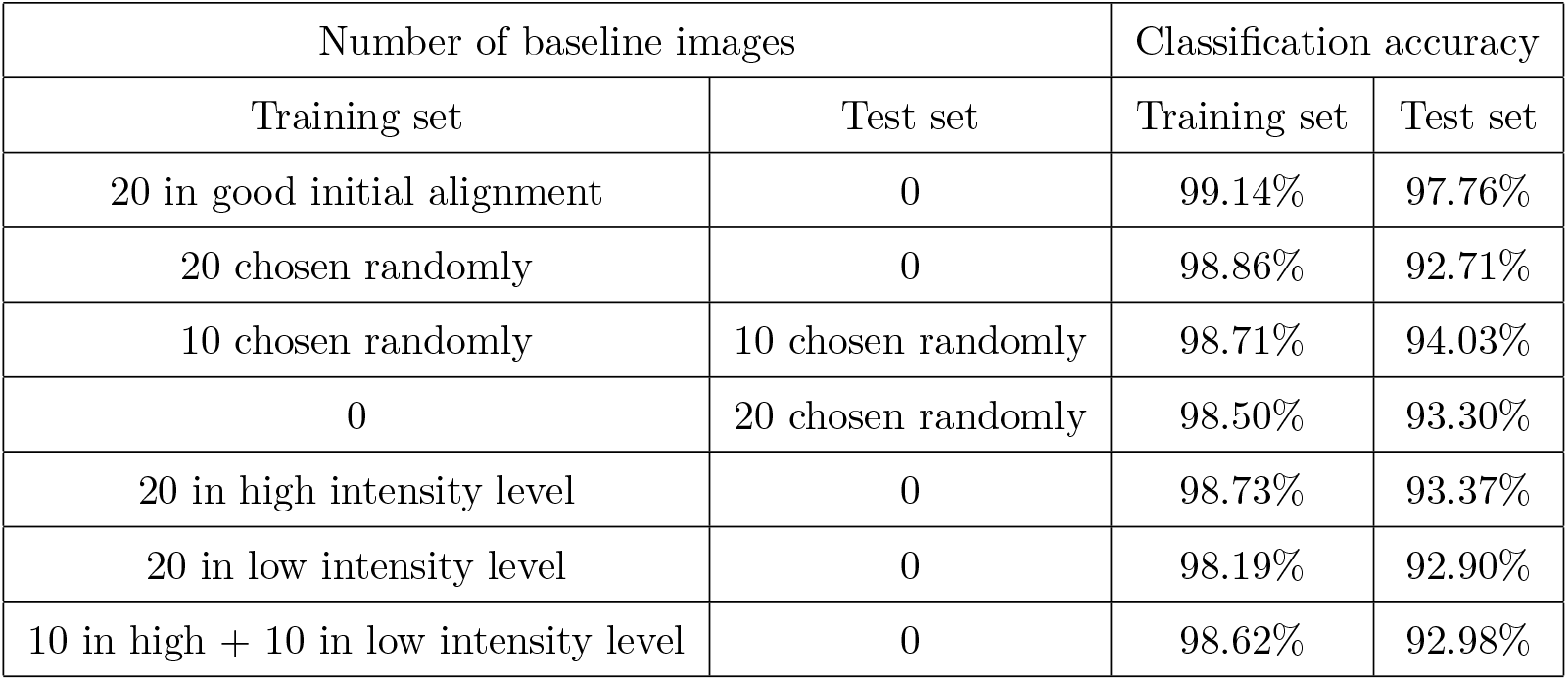
Seven distinct combinations of baseline images chosen to construct the PET template. For each experiment, all images were preprocessed using the constructed template followed by SIFT feature classification, and the accuracy with cross validation was recorded as a measure of the template quality.

| Number of baseline images |  | Classification accuracy |  |
| --- | --- | --- | --- |
| Training set | Test set | Training set | Test set |
| 20 in good initial alignment | 0 | 99.14% | 97.76% |
| 20 chosen randomly | 0 | 98.86% | 92.71% |
| 10 chosen randomly | 10 chosen randomly | 98.71% | 94.03% |
| 0 | 20 chosen randomly | 98.50% | 93.30% |
| 20 in high intensity level | 0 | 98.73% | 93.37% |
| 20 in low intensity level | 0 | 98.19% | 92.90% |
| 10 in high + 10 in low intensity level | 0 | 98.62% | 92.98% |

### 4.3. SIFT and feature pattern

The conventional SIFT algorithm locates key points by looking for the space extrema through the Difference of Gaussian (DoG). We tried to extract the space extrema for the 3D images in a four-dimensional neighborhood with the DoG. Only a few key points were returned from the calculation, while some images had no extrema. This was due to the nature of the PET images, where neighboring voxels were highly correlated because of the continuous FDG concentration in the brain and the smoothing step in the preprocessing. Thus, for our 3D PET images, which were registered to the template, we decided to use a fixed set of key points which are all in the regions that are shown in previous studies to be affected by the MCAO procedure. In this way, the extracted descriptors from different images had the same locations and thus could be compared among each other.

After concatenating the SIFT descriptors of 25 key points into one feature vector, we performed dimensionality reduction through PCA and obtained significant results. The two plots in Figure 4 both show a trend that the blue data points (normal images) tend to be clustered, whereas the orange points (MCAO images) have a larger variance. This phenomenon suggests that the MCAO surgery will have different influence on different rats due to individual differences or distinct trials [38]. The discrepancy of the MCAO outcome may lead to incorrect statistical decisions if we adopt the voxel-based analysis methods. By using the SVM model, we clearly show that such a discrepancy is still much smaller than the group-wise difference between normal images and MCAO images and can therefore be reliably treated as intra-class variation.

### 4.4. Limitations

Our study still has room for improvement. For example, there were thousands of voxels in the injured areas, so it was infeasible to check each of them one by one and find an optimal key point. We expect that researchers who are expert in knowing exactly where the brain ischemia and infarctions happen can provide more accurate key point locations. Additionally, the study of using other image features apart from SIFT is an interesting topic and an open problem.

## 5. Conclusion

This paper presented a protocol to assess the quality of rat MCAO procedure by extracting SIFT features and building an SVM model from the FDG microPET imaging data. It has advantages over behavior tests or TTC staining by providing quantitative and non-invasive measurements. Unlike existing voxel-based methods (such as the SPM) that suffer from inconsistent results due to hyper-FDG uptake or potential bias caused by uncontrollable FDG dosage difference, our protocol can provide consistent and reliable classification results without intensity normalization. Moreover, our method does not rely on comparison between scans before and after the MCAO, and therefore the baseline scan is not always required when designing the experiment. This will be beneficial when the proposed experiment involves large number of subjects.

## 6. Author contributions

Wuxian He: Methodology, software, formal analysis, writing - original draft

Hongtu Tang: Validation, investigation, Resources

Jia Li: Validation, investigation, Resources

Chenze Hou: Methodology, software

Xiaoyan Shen: Investigation, data curation

Chenrui Li: Software, data curation, visualization

Huafeng Liu: Conceptualization, project administration, funding acquisition

Weichuan Yu: Conceptualization, writing - review & editing, supervision, project administration, funding acquisition

## 8. Acknowledgements

We thank Jingwen Zheng for kindly helping setting up the webpage and Tamie KONSTAS for carefully proofreading the manuscript.

This work was supported in part by the National Natural Science Foundation of China (No: 61525106, U1809204), the National Key Technology Research and Development Program of China (No: 2017YFE0104000) and R4012-18 from the Research Grant Council (RGC) of the Hong Kong S.A.R. government of China.

## References

[1] W. Johnson, O. Onuma, M. Owolabi, S. Sachdev, Stroke: a global response is needed, Bulletin of the World Health Organization 94 (2016) 634. doi: 10.2471/BLT.16.181636.

[2] E. Z. Longa, P. R. Weinstein, S. Carlson, R. Cummins, Reversible middle cerebral artery occlusion without craniectomy in rats, Stroke 20 (1989) 84–91. doi:10.1161/01.str.20.1.84.

[3] J. B. Bederson, L. H. Pitts, M. Tsuji, M. Nishimura, R. Davis, H. Bartkowski, Rat middle cerebral artery occlusion: evaluation of the model and development of a neurologic examination, Stroke 17 (1986) 472–476. doi:10.1161/01.str.17.3.472.

[4] A. Benedek, K. Móricz, Z. Jurányi, G. Gigler, G. Lévay, L. G. Hársing Jr, P. Mátyus, G. Szenási, M. Albert, Use of TTC staining for the evaluation of tissue injury in the early phases of reperfusion after focal cerebral ischemia in rats, Brain Research 1116 (2006) 159–165. doi:10.1016/j.brainres.2006.07.123.

[5] M. Bieber, J. Gronewold, A.-C. Scharf, M. K. Schuhmann, F. Langhauser, S. Hopp, S. Mencl, E. Geuss, J. Leinweber, J. Guthmann, T. R. Doeppner, C. Kleinschnitz, G. Stoll, P. Kraft, D. M. Hermann, Validity and reliability of neurological scores in mice exposed to middle cerebral artery occlusion, Stroke 50 (2019) 2875–2882. doi:10.1161/STROKEAHA.119.026652.

[6] S. Ansari, H. Azari, D. J. McConnell, A. Afzal, J. Mocco, Intraluminal middle cerebral artery occlusion (MCAO) model for ischemic stroke with laser Doppler flowmetry guidance in mice, JoVE (Journal of Visualized Experiments) (2011) e2879 doi:10.3791/2879.

[7] E. Ingberg, H. Dock, E. Theodorsson, A. Theodorsson, J. O. Ström, Effect of laser Doppler flowmetry and occlusion time on outcome variability and mortality in rat middle cerebral artery occlusion: inconclusive results, BMC Neuroscience 19 (2018) 1–11. doi:10.1186/s12868-018-0425-0.

[8] V. S. Hedna, S. Ansari, S. Shahjouei, P. Y. Cai, A. S. Ahmad, J. Mocco, A. I. Qureshi, Validity of laser Doppler flowmetry in predicting outcome in murine intraluminal middle cerebral artery occlusion stroke, Journal of Vascular and Interventional Neurology 8 (2015) 74. doi:PMID:26301036.

[9] V. Rajan, B. Varghese, T. G. van Leeuwen, W. Steenbergen, Review of methodological developments in laser Doppler flowmetry, Lasers in Medical Science 24 (2009) 269–283. doi:https://doi.org/10.1007/s10103-007-0524-0.

[10] E. M. Lake, P. Bazzigaluppi, J. Mester, L. A. Thomason, R. Janik, M. Brown, J. McLaurin, P. L. Carlen, D. Corbett, G. J. Stanisz, B. Stefanovic, Neurovascular unit remodelling in the subacute stage of stroke recovery, NeuroImage 146 (2017) 869–882. doi:10.1016/j.neuroimage.2016.09.016.

[11] T. E. Lund, M. D. Nørgaard, E. Rostrup, J. B. Rowe, O. B. Paulson, Motion or activity: their role in intra-and inter-subject variation in fMRI, NeuroImage 26 (2005) 960–964. doi:10.1016/j.neuroimage.2005.02.021.

[12] J. D. Power, A. Mitra, T. O. Laumann, A. Z. Snyder, B. L. Schlaggar, S. E. Petersen, Methods to detect, characterize, and remove motion artifact in resting state fMRI, NeuroImage 84 (2014) 320–341. doi:10.1016/j.neuroimage.2013.08.048.

[13] H. Liu, X. Shen, H. Tang, J. Li, T. Xiang, W. Yu, Using microPET imaging in quantitative verification of the acupuncture effect in ischemia stroke treatment, Scientific Reports 3 (2013) 1–7. doi:10.1038/srep01070.

[14] W. C. Kreisl, M.-J. Kim, J. M. Coughlin, I. D. Henter, D. R. Owen, R. B. Innis, PET imaging of neuroinflammation in neurological disorders, The Lancet Neurology 19 (2020) 940–950. doi:10.1016/S1474-4422(20)30346-X.

[15] H. Barthel, V. Zeisig, B. Nitzsche, M. Patt, J. Patt, G. Becker, A. Dreyer, J. Boltze, O. Sabri, Cerebral blood flow measurement with oxygen-15 water positron emission tomography, in: PET and SPECT of Neurobiological Systems, 2021, pp. 127–152. doi:10.1007/978-3-642-42014-6_4.

[16] T. Maaniitty, J. Knuuti, A. Saraste, 15O-Water PET MPI: Current status and future perspectives, in: Seminars in Nuclear Medicine, 2020. doi: 10.1053/j.semnuclmed.2020.02.011.

[17] A. Bunevicius, H. Yuan, W. Lin, The potential roles of 18F-FDG-PET in management of acute stroke patients, BioMed Research International 2013 (2013). doi:10.1155/2013/634598.

[18] W. D. Penny, K. J. Friston, J. T. Ashburner, S. J. Kiebel, T. E. Nichols, Statistical parametric mapping: the analysis of functional brain images, Elsevier, 2011. doi:10.1016/B978-0-12-372560-8.X5000-1.

[19] K. J. Worsley, A. C. Evans, S. Marrett, P. Neelin, A three-dimensional statistical analysis for CBF activation studies in human brain, Journal of Cerebral Blood Flow & Metabolism 12 (1992) 900–918. doi:10.1038/jcbfm.1992.127.

[20] G. Flandin, K. J. Friston, Analysis of family-wise error rates in statistical parametric mapping using random field theory, Human Brain Mapping 40 (2019) 2052–2054. doi:10.1002/hbm.23839.

[21] M. Sobrado, M. Delgado, E. Fernández-Valle, L. García-García, M. Torres, J. Sánchez-Prieto, J. Vivancos, R. Manzanares, M. Moro, M. A. Pozo, I. Lizasoain, Longitudinal studies of ischemic penumbra by using 18F-FDG PET and MRI techniques in permanent and transient focal cerebral ischemiain rats, NeuroImage 57 (2011) 45–54. doi:10.1016/j.neuroimage.2011.04.045.

[22] H. Yuan, J. E. Frank, Y. Hong, H. An, C. Eldeniz, J. Nie, A. Bunevicius, D. Shen, W. Lin, Spatiotemporal uptake characteristics of [18] F-2-fluoro-2-deoxy-D-glucose in a rat middle cerebral artery occlusion model, Stroke 44 (2013) 2292–2299. doi:10.1161/STROKEAHA.113.000903.

[23] F. J. López-González, J. Silva-Rodríguez, J. Paredes-Pacheco, A. Niñerola-Baizáan, N. Efthimiou, C. Martín-Martín, A. Moscoso, Á. Ruibal, N. Roé-Vellváe, P. Aguiar, Intensity normalization methods in brain FDG-PET quantification, NeuroImage 222 (2020) 117229. doi:10.1016/j.neuroimage.2020.117229.

[24] B. Nie, S. Liang, X. Jiang, S. Duan, Q. Huang, T. Zhang, P. Li, H. Liu, B. Shan, An automatic method for generating an unbiased intensity normalizing factor in positron emission tomography image analysis after stroke, Neuroscience Bulletin 34 (2018) 833–841. doi:10.1007/s12264-018-0240-8.

[25] B. Rister, D. Reiter, H. Zhang, D. Volz, Scale-and orientation-invariant keypoints in higher-dimensional data, in: 2015 IEEE International Conference on Image Processing (ICIP), 2015, pp. 3490–3494. doi:10.1109/ICIP.2015.7351453.

[26] D. G. Lowe, Distinctive image features from scale-invariant keypoints, International Journal of Computer Vision 60 (2004) 91–110. doi:10.1023/B:VISI.0000029664.99615.94.

[27] Y.-Y. Li, B. Zhang, K.-W. Yu, C. Li, H.-Y. Xie, W.-Q. Bao, Y.-Y. Kong, F.-Y. Jiao, Y.-H. Guan, Y.-L. Bai, Effects of constraint-induced movement therapy on brain glucose metabolism in a rat model of cerebral ischemia: a micro PET/CT study, International Journal of Neuroscience 128 (2018) 736–745. doi:10.1080/00207454.2017.1418343.

[28] P. Schweinhardt, P. Fransson, L. Olson, C. Spenger, J. L. Andersson, A template for spatial normalisation of MR images of the rat brain, Journal of Neuroscience Methods 129 (2003) 105–113. doi:10.1016/s0165-0270(03)00192-4.

[29] K. Vunckx, P. Dupont, K. Goffin, W. Van Paesschen, K. Van Laere, J. Nuyts, Voxel-based comparison of state-of-the-art reconstruction algorithms for 18F-FDG PET brain imaging using simulated and clinical data, NeuroImage 102 (2014) 875–884. doi:10.1016/j.neuroimage.2014.06.068.

[30] A. Popp, N. Jaenisch, O. W. Witte, C. Frahm, Identification of ischemic regions in a rat model of stroke, PloS One 4 (2009) e4764. doi:10.1371/journal.pone.0004764.

[31] E. A. Papp, T. B. Leergaard, E. Calabrese, G. A. Johnson, J. G. Bjaalie, Waxholm Space atlas of the Sprague Dawley rat brain, NeuroImage 97 (2014) 374–386. doi:10.1016/j.neuroimage.2014.04.001.

[32] J. Ahrens, B. Geveci, C. Law, Paraview: An end-user tool for large data visualization, The Visualization Handbook 717 (2005). doi:10.1016/B978-012387582-2/50038-1.

[33] P. Baumgartner, M. El Amki, O. Bracko, A. R. Luft, S. Wegener, Sensorimotor stroke alters hippocampo-thalamic network activity, Scientific Reports 8 (2018) 1–11. doi:10.1038/s41598-018-34002-9.

[34] T. Hastie, R. Tibshirani, J. Friedman, The elements of statistical learning: data mining, inference, and prediction, Springer Science & Business Media, 2009. doi:10.1007/978-0-387-84858-7.

[35] K. J. Worsley, S. Marrett, P. Neelin, A. C. Vandal, K. J. Friston, A. C. Evans, A unified statistical approach for determining significant signals in images of cerebral activation, Human Brain Mapping 4 (1996) 58–73. doi:10.1002/(SICI)1097-0193(1996)4:1<58::AID-HBM4>3.0.CO;2-0.

[36] C.-y. O. Wong, J. Thie, M. Gaskill, R. Ponto, J. Hill, H.-y. Tian, H. Balon, D. Wu, D. Fink-Bennett, C. Nagle, A statistical investigation of normal regional intra-subject heterogeneity of brain metabolism and perfusion by F-18 FDG and O-15 H_2_O PET imaging, BMC Nuclear Medicine 6 (2006) 4. doi:10.1186/1471-2385-6-4.

[37] J. Ashburner, K. J. Friston, W. Penny, Chapter 37-the general linear model, in: Human Brain Function (Second Edition), Academic Press, 2004, pp. 725–760. doi:10.1016/B978-012264841-0/50039-1.

[38] C. Leithner, M. Füchtemeier, D. Jorks, S. Mueller, U. Dirnagl, G. Royl, Infarct volume prediction by early magnetic resonance imaging in a murine stroke model depends on ischemia duration and time of imaging, Stroke 46 (2015) 3249–3259. doi:10.1161/STROKEAHA.114.007832.

