## Supplemental materials for "Feature-based Quality Assessment of Middle Cerebral Artery Occlusion Using 18F-fluorodeoxyglucose Positron Emission Tomography"

Image-based Quality Assessment of Middle Cerebral Artery Occlusion using SIFT Descriptor and Support Vector Machine

*Contributed equally

**Supplemental methods**

***Animals***

All the animal experiments in this study were approved by the Animal Research Committee of School of Medicine, Zhejiang University. The subjects involved in this study were adult female Sprague-Dawley rats with a weight of 180-280 g. Two independent middle cerebral artery occlusion (MCAO) surgeries were performed following the same procedure,[1] one with 116 rats and the other with 45 rats. Focal cerebral ischemia was induced through right brain MCAO: The squamosal and zygomatic bones in the right brain were exposed with skin incision and muscle excarnation. A high speed drill was used to make a small hole in the junction part of squamosal and zygomatic bones. Then the right middle cerebral artery was uncovered and occluded with bipolar electrical coagulation permanently. During the experiment, each rat had free access to food and water, and was anesthetized using 400 mg/kg chloral hydrate before the scanning. The entire experimental procedure followed the guidelines for the care and use of laboratory animals published by the National Institute of Health. All the data are available from the website: http://yulab.ust.hk/MCAO.

***Construction of rat FDG-PET template***

To transform PET scans of distinct rats into a standard space, a template is necessary. Since there is no widely accepted PET template for Sprague-Dawley rats, we constructed our own template from an existing MRI template1[2] using SPM12 (Wellcome Department of Cognitive Neurology, London, UK). The detailed steps are as follows:

(1) Choose 20 raw images of resting state (baseline) from the training set.
(2) Adjust the voxel size of the images to [9 9 14.4]. In this way, the rat brain can approximately fit the human brain size in order to use the default settings of SPM.
(3) Get all the images into a standard orientation as shown in Figure 1 below.
(4) Manually align the 20 images to the MRI template in SPM. This step aims to ensure the accuracy of the next normalization step.
(5) One of the 20 images is selected, and all the remaining ones are spatially normalized to it using SPM normalization function. The bounding box should be carefully set so that all the intracranial parts are included.
(6) The 20 spatially normalized images are averaged to create a mean image. Then co-register it to the MRI template in SPM to create the final FDG-PET template for rat brain.

The constructed FDG-PET template is provided on our website for free download and research.

***Preprocessing of PET images***

The raw images after reconstruction were preprocessed with the constructed template in SPM12. Detailed steps are shown as follows:

(1) The same as step (2) in the last section for the template construction.
(2) The same as step (3) in the last section.
(3) One image is selected, and all other images are aligned to it using SPM realignment function.
(4) Then the selected image is manually aligned to the PET template so that the

intracranial parts are roughly registered. The transformation parameters are applied to all remaining images so that they are aligned to the template as well.

(5) After that, all the images are co-registered to the PET template using SPM co-registration function (estimate only).

(6) Use the parameters from the previous step to reslice the images into the space defined by the ready-to-use image called target.img. The bounding box of the target image is slightly larger than the PET template, and the purpose is to leave more margin in the next normalization step. The target image can be downloaded from the website while people can also make it themselves.

(7) (For scans after MCAO only) The injured right hemisphere is replaced by the left hemisphere to produce a synthetic and symmetric image. The reason to do so is because injured areas will contribute much to the cost function during spatial normalization at the cost of matching accuracy in the healthy areas.

(8) The synthetic image is spatially normalized to the space defined by the PET template using SPM normalization function (estimate only) and the parameters are applied to the original image. For normal (baseline) images taken before MCAO, normalize the results from step 6 to the PET template. The bounding box is set as [-78 -154 -118; 80 60 6].

(9) Remove extracranial tissues by a ready-to-use mask image. It is a binary image and people can make it by segmentation of intracranial tissues from the template. We also provide it on the website.

(10) All images are smoothed using a 16 mm isotropic Gaussian kernel.

***3D scale invariant feature transform (SIFT)***

After spatial preprocessing, image features were extracted using 3D SIFT method.[3] The Original SIFT is used for 2D images and the detailed algorithm can be found in the original paper by Lowe.[4] The 2D SIFT briefly has four steps:

(1) The image is convolved using Gaussian functions with distinct variance, which results in a scale space (stacks of Gaussian images with different scales). Then, calculate the Difference of Gaussian (DoG) by subtracting adjacent Gaussian images. Detect the minima or maxima of the DoG images by comparing each pixel with its 26 neighbors in the current and its two adjacent DoG images. The computed extreme pixels are regarded as candidate key points.

(2) There will be redundant and unstable candidate key points detected from the previous step. After interpolation of nearby pixels, reject key points that have low contrast or unstable edge responses. Thus, the location of scale and space invariant key points are finally determined.

(3) Define the gradient and orientation for each key point. For an image L(x, y), its gradient magnitude m(x, y) and orientation Θ(x; y) are defined as below:

$$m(x,y) = \sqrt{\left(\left(L(x+1,y) - L(x-1,y)\right)^2 + \left(L(x,y+1) - L(x,y-1)\right)^2\right)}$$

$$\theta(x,y) = \tan^{-1}\left(\left(L(x,y+1) - L(x,y-1)\right)/\left(L(x+1,y) - L(x-1,y)\right)\right)$$

(4) Obtain a descriptive vector for each key point such that it will be highly distinctive

and invariant to location, scaling, translation, rotation and illumination. The descriptor of a key point is constructed from the gradient magnitude and orientation for each sample point in the neighborhood of the key point.

Since PET images were three dimensional, we adopted a 3D SIFT algorithm to extract features. This algorithm extends the notions from 2D SIFT to 3D by using the icosahedral histograms with 12 vertices. In conventional 2D SIFT, the gradient has 8 directions which comes from dividing the planar circle into 8 sectors equally. In the 3D scenario, this algorithm tessellates the sphere into 12 equal tiles. Then the length of a SIFT descriptor will be lifted from 128 to 768. Detailed algorithm of this 3D SIFT can be found in Rister et al.[3]

The coordinate information of the 25 key points is provided on the website. The 3D rendered figure for visualization using ParaView[5] is shown in Figure 2 below. The 25 key points were chosen from brain regions affected by stroke,[6] and the SIFT descriptor of each key point is calculated. Then, the 25 descriptors were concatenated into a single vector, which became the image feature. Brain region delineation is based on the Waxholm atlas (3rd version),[7] and the names of the 118 defined regions are shown in Table 1. The original resolution of the atlas is different from our PET template, so it needs to be normalized to the template using SPM normalization function. Then the 3D coordinates of voxels in each brain area were looked up by the intensity values provided from the atlas.

The 3D SIFT descriptor is a 768-component vector for each key point. Since each image has 25 key points, we can concatenate the descriptors into a long feature vector with 768*25=19200 components. The feature vectors will be used in the SVM classification for MCAO quality assessment.

**Supplemental Tables**

Supplemental Table 1. List of brain regions and their intensity values in the atlas. Region names in **BOLD** are those where we extracted key points.

| *No.* | Region name | Value | *No.* | Region name | Value |
|---|---|---|---|---|---|
| *1* | Descending corticofugal pathways | 1 | *60* | Supraoptic decussation | 83 |
| *2* | Substantia nigra | 2 | *61* | Medial lemniscus decussation | 84 |
| *3* | **Subthalamic nucleus** | 3 | *62* | Pyramidal decussation | 85 |
| *4* | Molecular layer of the cerebellum | 4 | *63* | **Neocortex** | 92 |
| *5* | Granule cell level of the cerebellum | 5 | *64* | Bed nucleus of the stria terminalis | 93 |
| *6* | **Alveus of the hippocampus** | 6 | *65* | Pretectal region | 94 |
| *7* | Inferior cerebellar peduncle | 7 | *66* | Cornu ammonis 3 | 95 |
| *8* | Cingulate cortex, area 2 | 10 | *67* | Dentate gyrus | 96 |
| *9* | **Striatum** | 30 | *68* | Cornu ammonis 2 | 97 |
| *10* | Globus pallidus | 31 | *69* | Cornu ammonis 1 | 98 |
| *11* | Entopeduncular nucleus | 32 | *70* | Fasciola cinereum | 99 |
| *12* | Ventricular system | 33 | *71* | Subiculum | 100 |
| *13* | Medial lemniscus | 34 | *72* | **Postrhinal cortex** | 108 |
| *14* | Facial nerve | 35 | *73* | Presubiculum | 109 |
| *15* | Anterior commissure, anterior part | 36 | *74* | Parasubiculum | 110 |
| *16* | Anterior commissure, posterior part | 37 | *75* | **Perirhinal area 35** | 112 |
| *17* | **Ventral hippocampal commissure** | 38 | *76* | **Perirhinal area 36** | 113 |
| *18* | **Thalamus** | 39 | *77* | Entorhinal cortex | 114 |
| *19* | Septal region | 40 | *78* | Lateral entorhinal cortex | 115 |
| *20* | Optic nerve | 41 | *79* | Vestibular apparatus | 119 |
| *21* | Optic tract and optic chiasm | 42 | *80* | Cochlea | 120 |
| *22* | Pineal gland | 43 | *81* | Cochlear nerve | 121 |
| *23* | Spinal cord | 45 | *82* | Vestibular nerve | 122 |
| *24* | Commissure of the superior colliculus | 46 | *83* | Ventral cochlear nucleus, granule cell layer | 123 |
| *25* | Brainstem | 47 | *84* | 4th ventricle | 125 |
| *26* | Hypothalamic region | 48 | *85* | Dorsal cochlear nucleus, molecular layer | 126 |
| *27* | Superficial gray layer of the superior colliculus | 50 | *86* | Dorsal cochlear nucleus, fusiform and granule layer | 127 |

| | | | | | |
|---|---|---|---|---|---|
| *28* | Periaqueductal gray | 51 | *87* | Dorsal cochlear nucleus, deep core | 128 |
| *29* | Fornix | 52 | *88* | Acoustic striae | 129 |
| *30* | Mammillothalamic tract | 53 | *89* | Trapezoid body | 130 |
| *31* | Commissural stria terminalis | 54 | *90* | Nucleus of the trapezoid body | 131 |
| *32* | Deeper layers of the superior colliculus | 55 | *91* | Superior paraolivary nucleus | 132 |
| *33* | Periventricular gray | 56 | *92* | Medial superior olive | 133 |
| *34* | Genu of the facial nerve | 57 | *93* | Lateral superior olive | 134 |
| *35* | Pontine nuclei | 58 | *94* | Superior periolivary region | 135 |
| *36* | **Fimbria of the hippocampus** | 59 | *95* | Ventral periolivary nuclei | 136 |
| *37* | Fasciculus retroflexus | 60 | *96* | Lateral lemniscus, ventral nucleus | 137 |
| *38* | **Stria medullaris of the thalamus** | 61 | *97* | Lateral lemniscus, intermediate nucleus | 138 |
| *39* | Stria terminalis | 62 | *98* | Lateral lemniscus, dorsal nucleus | 139 |
| *40* | Posterior commissure | 63 | *99* | Lateral lemniscus, commissure | 140 |
| *41* | Glomerular layer of the accessory olfactory bulb | 64 | *100* | Lateral lemniscus | 141 |
| *42* | Glomerular layer of the olfactory bulb | 65 | *101* | Inferior colliculus, dorsal cortex | 142 |
| *43* | Olfactory bulb | 66 | *102* | Inferior colliculus, central nucleus | 143 |
| *44* | Corpus callosum and associated subcortical white matter | 67 | *103* | Inferior colliculus, external cortex | 145 |
| *45* | Brachium of the superior colliculus | 68 | *104* | Inferior colliculus, brachium | 146 |
| *46* | Inferior colliculus, commissure | 69 | *105* | Medial geniculate body, medial division | 147 |
| *47* | Central canal | 70 | *106* | Medial geniculate body, dorsal division | 148 |
| *48* | **Interpeduncular nucleus** | 71 | *107* | Medial geniculate body, ventral division | 149 |
| *49* | Ascending fibers of the facial nerve | 72 | *108* | Medial geniculate body, marginal zone | 150 |
| *50* | Anterior commissure | 73 | *109* | Primary auditory cortex | 151 |
| *51* | Inferior olive | 74 | *110* | Secondary auditory cortex, dorsal area | 152 |
| *52* | Spinal trigeminal nucleus | 75 | *111* | Secondary auditory cortex, ventral area | 153 |
| *53* | Spinal trigeminal tract | 76 | *112* | Auditory radiation | 157 |

| | | | | | |
|---|---|---|---|---|---|
| *54* | Frontal association cortex | 77 | *113* | ventral cochlear nucleus, anterior part | 158 |
| *55* | **Middle cerebellar peduncle** | 78 | *114* | Ventral cochlear nucleus, posterior part | 159 |
| *56* | Transverse fibers of the pons | 79 | *115* | Ventral cochlear nucleus, cap area | 160 |
| *57* | Habenular commissure | 80 | *116* | Spiral ganglion | 162 |
| *58* | Nucleus of the stria medullaris | 81 | *117* | Nucleus sagulum | 163 |
| *59* | Basal forebrain region | 82 | *118* | Reticular thalamic nucleus, auditory segment | 164 |

**Supplemental Figures**

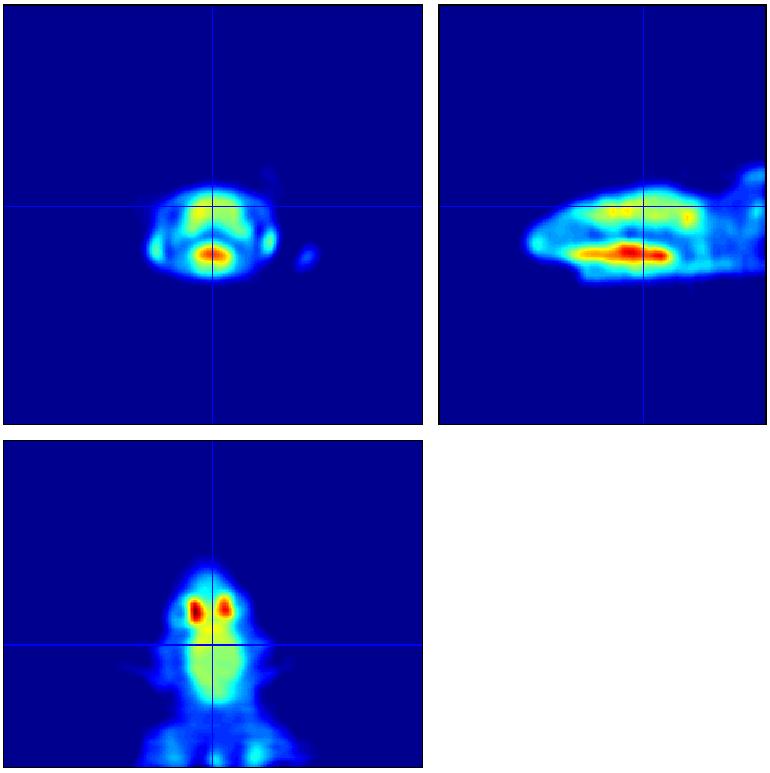

**Supplemental Figure 1.** An example of raw image in the standard orientation with proper voxel size.

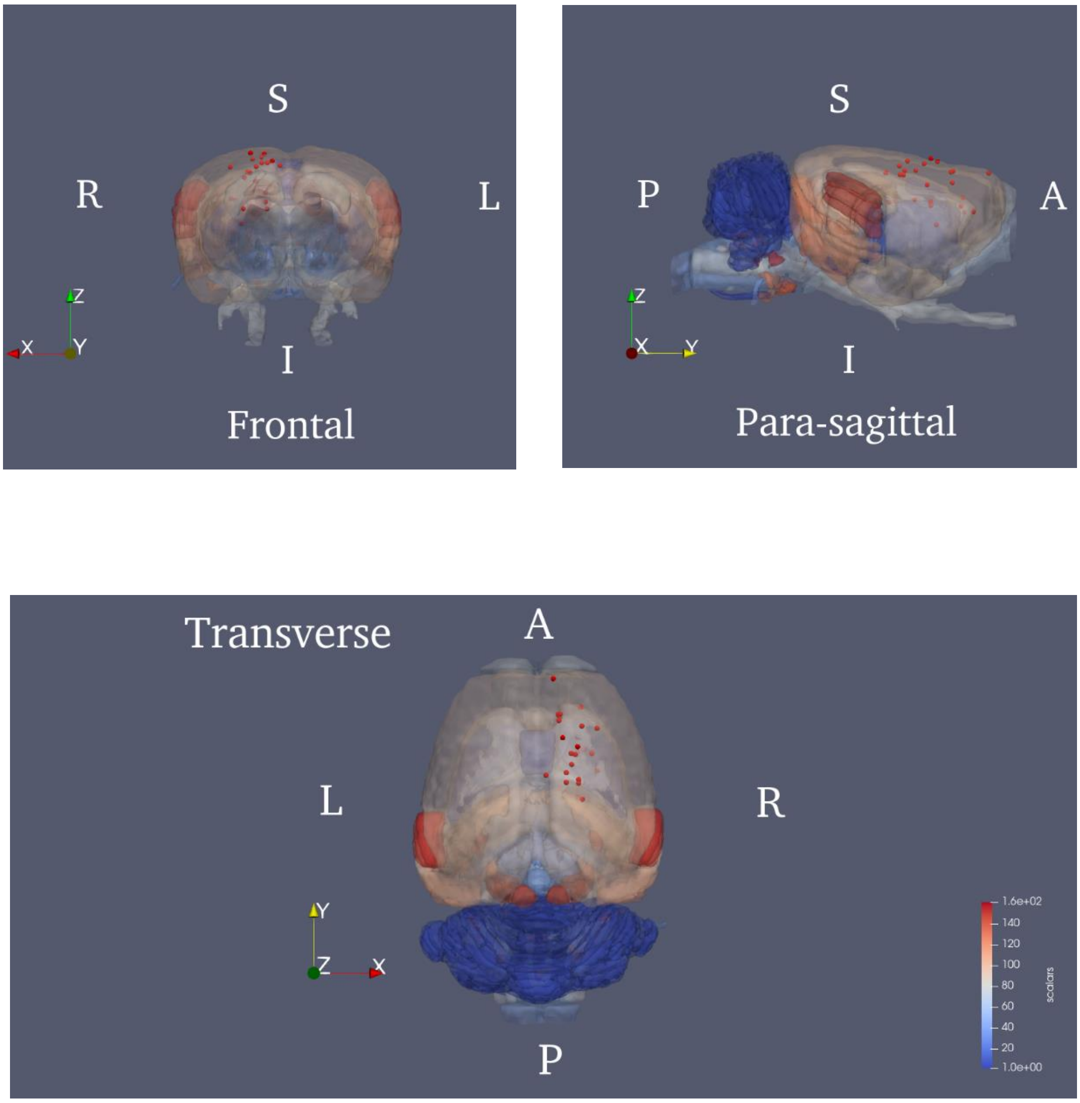


**Supplemental Figure 2.** The 3D rendered atlas with the 25 chosen key points (all from the right hemisphere) to extract the image features. (R, right; L, left; S, superior; I, inferior; P, posterior; A, anterior)

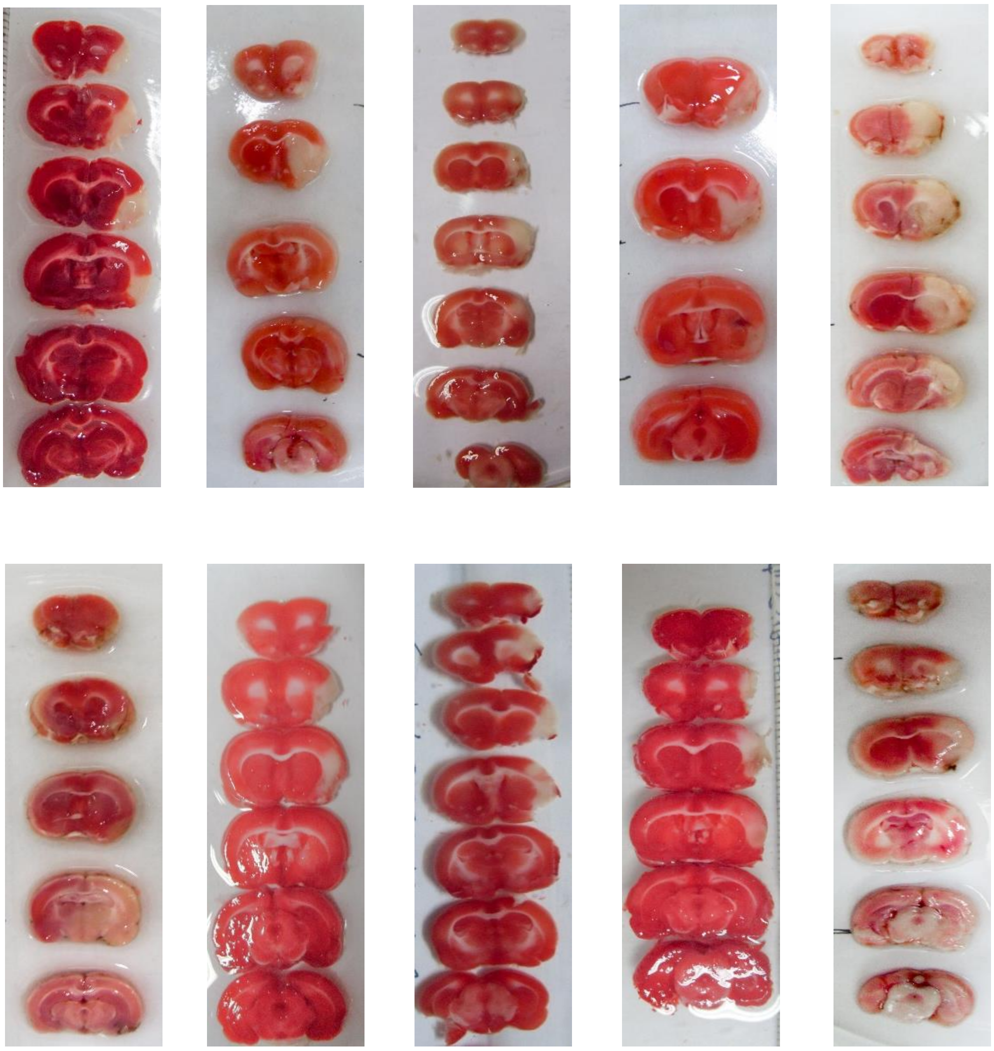

**Supplemental Figure 3.** Triphenyl tetrazolium chloride (TTC) staining results of the 10 rats in the validation set.

**Supplemental References**


1. Longa EZ, Weinstein PR, Carlson S, Cummins R. Reversible middle cerebral artery occlusion without craniectomy in rats. *Stroke*. 1989;20:84-91.
2. Schweinhardt P, Fransson P, Olson L, Spenger C, Andersson J L. A template for spatial normalisation of MR images of the rat brain. Journal of Neuroscience Methods. 2003;129:105-113
3. Rister B, Reiter D, Zhang H, Volz D, Horowitz M, Gabr RE, Cavallaro JR. Scale- and orientation-invariant keypoints in higher-dimensional data. Paper presented at: 2015 IEEE International Conference on Image Processing (ICIP); September 27-30, 2015; Quebec City, Canada. (pp. 3490-3494). IEEE. https://ieeexplore.ieee.org/abstract/document/7351453. Accessed November 9, 2020.
4. Lowe DG. Distinctive image features from scale-invariant keypoints. International Journal of Computer Vision. 2004;60:91-110.
5. Ahrens J, Geveci B, Law C. Paraview: An end-user tool for large data visualization. The Visualization Handbook. Elsevier; 2005.
6. Popp A, Jaenisch N, Witte OW, Frahm C. Identification of ischemic regions in a rat model of stroke. PloS one. 2009;4:e4764.
7. Papp EA, Leergaard TB, Calabrese E, Johnson GA, Bjaalie JG. Waxholm Space atlas of the Sprague Dawley rat brain. Neuroimage. 2014;97:374-386.
